# Prevalence of Depression among Epileptic Patients and its Association with Drug Therapy: A Systematic Review and Meta-Analysis

**DOI:** 10.1101/387571

**Authors:** Getenet Dessie, Henok Mulugeta, Cheru Tessema Leshargie, Fasil Wagnew, Sahai Burrowes

**Affiliations:** Lecturer of Nursing, Department of Nursing, College of Health Science Debre Markos University, Debre Markos Ethiopia Address: P.O. Box 269, Debre Markos, Ethiopia; Lecturer of Environmental Health, Department of Environmental Health, College of Health Science Debre Markos University, Debre Markos Ethiopia Address: P.O. Box 269, Debre Markos, Ethiopia; Public Health Program, College of Education and Health Sciences, Touro University, California 1310 Club Dr. Mare Island, Vallejo, CA 94592

**Keywords:** depression, drug therapy, epilepsy, prevalence, meta-analysis, Africa

## Abstract

**Background:** Despite the high prevalence of epilepsy in sub-Saharan Africa and the established relationship between depression and epilepsy, the extent of comorbid epilepsy and depression in the region is still poorly understood. This systematic review and meta-analysis aims to address this gap in the literature by determining the pooled prevalence of depression among epileptic patients in sub-Saharan Africa.

**Methods:** A systematic desk review and electronic web-based search of PubMed, Google Scholar, EMBASE, PsycINFO and the World Health Organization’s Hinari portal (which includes the SCOPUS, African Index Medicus, and African Journals Online databases) identified peer-reviewed research studies and doctoral dissertations on the prevalence of depression among patients with epilepsy using pre-defined quality and inclusion criteria. Relevant data were extracted and descriptive summaries of the studies presented in tabular form. The I^2^ test was used to assess heterogeneity across studies. A random effects model was used to estimate the pooled prevalence of comorbidity at a 95% confidence interval (CI). Funnel plot asymmetry and Egger’s tests were used to check for publication bias. The final effect size was determined by applying a trim and fill analysis in a random-effects model.

**Results:** Our search identified 167 studies, of which 16 articles were eligible for inclusion the final analysis. The pooled estimate of prevalence of depression among patients with epilepsy was 32.71 (95% CI: 25.50 - 39.91). Regional sub-group analysis found that the pooled prevalence in East Africa was 34.52 (95% CI: 23.53 - 45.51) and 29.69 (95% CI: 22.7 - 36.68) in Southern and West Africa. The odds of depression among epileptic patients receiving polytherapy were 2.65 higher than in those receiving monotherapy (95% CI: 1.49 - 4.71, I^2^=79.1%, p < 0.05).

**Conclusion:** Our findings indicate high comorbidity in sub-Saharan Africa and suggests that it may be more prevalent there than elsewhere. Comorbidity is statistically associated with polytherapy. Given the high levels of epilepsy in the region, more attention should be paid to incorporating depression screening and treatment into existing epilepsy programs and to revising treatment guidelines on comorbid depression to reduce polytherapy.

## Introduction

Epilepsy is one of the most common neurological disorders globally and in sub-Saharan Africa where it affects approximately 10 million people annually(1). Untreated, epilepsy can cause traumatic injuries, impair physical functioning, and reduce social engagement. This in turn can result in significant psychological stress and premature death.

People with epilepsy (PWE) are more vulnerable to psychiatric illnesses. Rates of psychiatric illness are 9% higher among PWE that in the general population and rates of depression, 22% higher. Depression is the most common psychiatric disorder in PWE(2) and major depressive episodes are one of the most common diagnoses among PWE. Social stigma, feelings of frustration and low self-esteem due to the danger and unpredictability of the illness, and the psychotropic effects of antiepileptic drugs (AEDs) have all been posited as reasons for the strong association between the two illnesses(3). There is also evidence that the association between depression and epilepsy may be bi-directional with people who are depressed having higher risk for developing epilepsy perhaps due to higher rates of head injuries and substance abuse(2)

Having a depression-related comorbidity is associated with in poorer quality of life and increased suicidal ideation for epileptic patients (4, 5). Greater severity of comorbid depression with epilepsy is associated with significantly reduced overall seizure recovery, higher seizure severity, and increased cognitive, emotional, and physical illness (6, 7). Clinicians may also find managing anti-depression treatment particularly challenging for their patients with epilepsy due to concerns about drug interactions, the side effects of polytherapy, and fears of lowering seizure thresholds (8, 9).

In addition to complicating treatment and reducing the quality of life, depression in PWE also places a strain on health systems, particularly in low-income countries because patients with untreated depression tend to use significantly more health resources. For example, Cramer et al’s study of health care utilization among comorbid patients found that epileptic patients with mild to moderate depression had a two-fold increase in medical visits, and those with severe depression, a four-fold increase, compared to those who were not depressed (6, 10). Moreover, studies suggest that depressed patients might be less adherent to epilepsy treatment than their non-depressed counterparts (11)and may respond relatively poorly to drug treatment(12).

Several factors have been found to be associated with increased risk of depression in the epileptic population. In sub-Saharan Africa, lower educational status, lower monthly income, frequency of seizure, the side effects of AEDs, and difficulties adhering to AEDs have all been found to be prominent risk factors for depression (13–17). Polytherapy has also been reported as an important factor but the association between depression and polypharmacy exhibits significant variation across studies; some finding a significant positive association (13–15, 18) and others none (16).

Studying the psychiatric comorbidities of epilepsy in sub-Saharan Africa is important because of the high prevalence of epilepsy in the region. This elevated prevalence is thought to be due to the endemicity of bacterial and parasitic infections that affect the central nervous system and to poor labor and delivery and perinatal care practices that could result in head trauma in infants and young children (19, 20) It is also reasonable to expect that the prevalence of depression comorbidity and its negative health and socio-economic effects would be more pronounced in the sub-Saharan African region where social stigma surrounding epilepsy is more pronounced, and the availability of adequate treatment lacking. However, studying the extent of depression and epilepsy comorbidity in sub-Saharan Africa is complicated by the fact that only about 20% of PWE in low- and middle-income countries receive treatment(1) and by poor estimates of the underlying population prevalence of both diseases. While there is relatively consistent evidence of high prevalence of comorbidity globally, most systematic reviews to date have included either no African studies (21, 22) or only one or two of studies from the continent (23) while the African literature on the subject has been characterized by considerable variability, inconsistency, and inconclusive findings.

Better information on the extent of this comorbidity is important for reducing inappropriate treatment and for improving suicide prevention in this vulnerable population (24). This systematic review and meta-analysis therefore, aims to synthesize evidence on the prevalence of depression among epileptic adults and adolescents and its association with drug therapy in Sub-Saharan Africa.

## Methods

### Search approach and appraisal of studies

Articles reviewed in this meta-analysis were accessed through electronic web-based database searches, desk reviews of the grey literature, and reference list reviews using the Preferred Reporting Items of Systematic Reviews and Meta-Analysis (PRISMA) checklist guidelines(25). The electronic databases searched were PubMed, Google Scholar, embase, PsycINFO and a World Health Organization (WHO) database portal for low- and middle-income countries that includes the Web of Science, SCOPUS, African Index Medicus (AIM), Cumulative Index to Nursing and Allied Health Literature (CINAHL), WHO’s Institutional Repository for Information Sharing (IRIS) and African Journals Online databases. In addition, the researchers found related articles through a desk review of the grey literature available on local shelves grey literature and from reviewing the reference lists of related articles.

The researchers used the following key terms for the database searches: *“*depression*” AND “*epilepsy*” OR “*co-morbid depression*” AND “*epilepsy*” OR “mental illness” AND “*sub-Saharan Africa*”.* Searches were conducted from December 1, 2017 to January 30, 2018.

### Inclusion and exclusion criteria

All English-language, full-text articles on observational studies (case-control or cross sectional) conducted in the sub-Saharan Africa region from 2005 to 2017, with adults or adolescents that were published in peer-reviewed journals or grey literature, who use internationally accepted scales to measure epilepsy and depression, who define epilepsy and depression according to internationally accepted definition and that concerned comorbid depression and epilepsy were eligible for inclusion.

#### Data abstraction and quality assessment

After initial screening, two reviewers (FW and GD) downloaded abstracts to assess them for inclusion. The relevance of the articles was evaluated based on their topic, objectives, and methodology as listed in the abstract. The abstracts were also assessed for agreement with the inclusion criteria. When it was unclear whether an abstract was relevant, it was included for retrieval. At this stage articles deemed irrelevant or out of the scope of the study were excluded and the full text of the remainder downloaded for a detailed review. Two reviewers (GD and FW) then assessed the quality of potentially eligible articles using the Newcastle-Ottawa Scale (NOS) criteria (26). Discrepancies in quality assessment scores were resolved with a third reviewer (HM), whenever appropriate. The average of two independent reviewers’ score was used to determine whether the articles should be included. Articles whose NOS quality scores were less than six, or that had methodological flaws, incomplete reporting of results, or for which full text was not available, were excluded from the final analysis. Study researchers made two separate attempts to contact article authors whenever additional study information was needed; for example, when patient outcome data were incomplete.

#### Outcome of interest

The outcome of interest was the pooled prevalence of depression among epileptic patients in sub-Saharan Africa. The prevalence is measured as the number of comorbid study subjects divided by the number of patients in a study multiplied by 100. We also estimate the association (as measured by crude odds ratios) between comorbidity and polypharmacy as a secondary outcome.

#### Data analysis

Information on the study characteristics (time frame, study location, study design, sample size, number of comorbid patients, and the age-range of patients) was extracted from each study using a Microsoft Excel spreadsheet template. These data were then transferred to Stata version 11 software to describe the pooled prevalence of depression among epileptic patients and to identify the significant association between the outcome variable and factors. The heterogeneity of study outcomes was assessed using the I^2^ test (27) and the Cochran’s Q-statistic. Because the test statistic indicated significant heterogeneity among studies (I^2^ >70%, p<0.05) a random effects model was used to estimate the pooled prevalence of comorbidity at a 95% confidence interval (CI) and a geographic subgroup analysis was conducted. We used funnel plot asymmetry (Figure 1) and Egger’s and Begg-Mazumdar Rank correlation tests to check for publication bias (28). Because the results of these tests suggested the possible existence of significant publication bias, the final effect size was determined by applying trim and fill analysis in the random-effects model (29). To confirm results, two researchers independently carried out the main statistical analysis and results were cross-checked for consistency.

**Figure 1.**
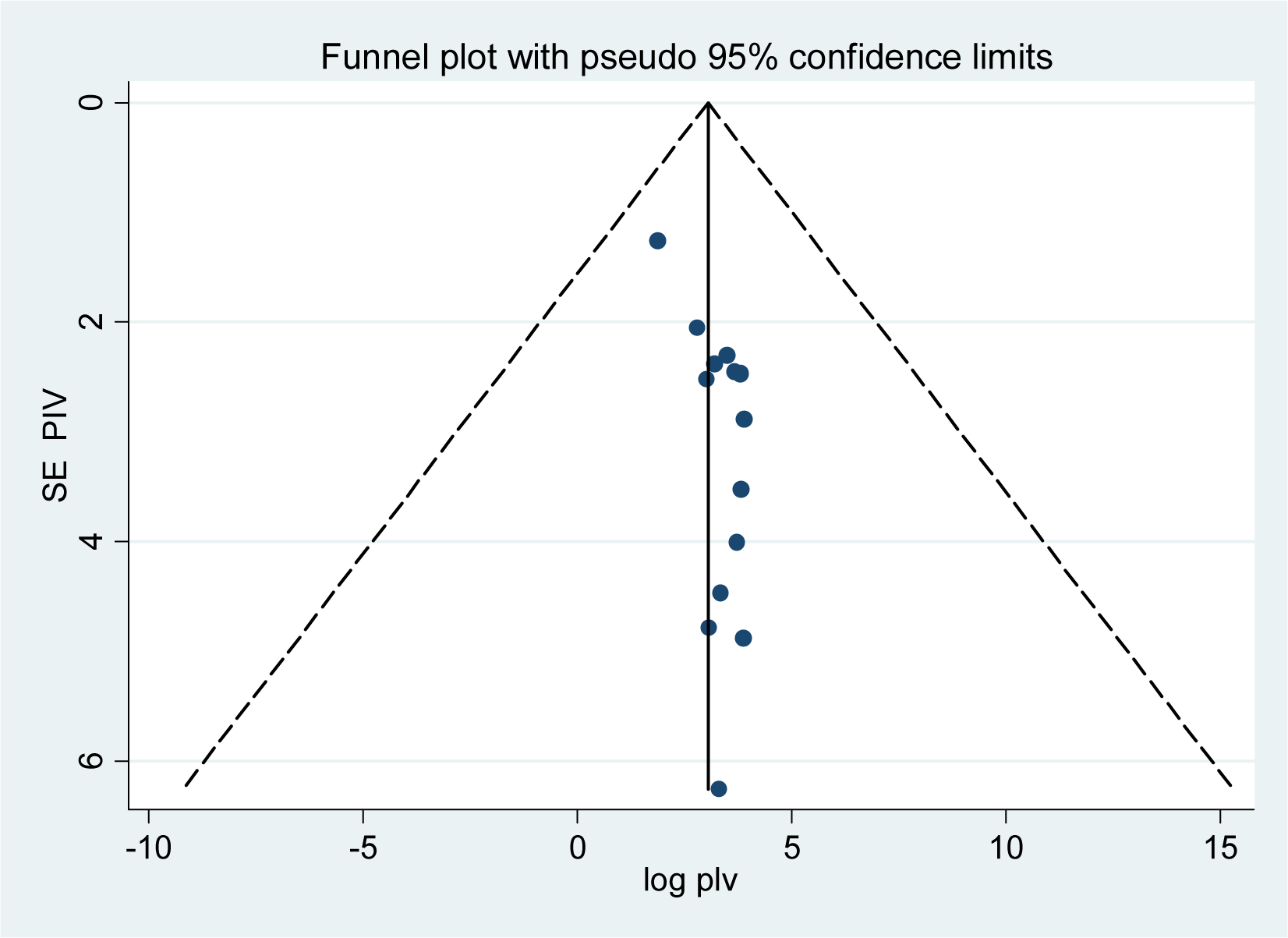
Meta funnel presentation of the prevalence of depression among epileptic patients, **2005–2017,** Saharan Africa.

## Results

### Identification and description of studies

The database search and desk review yielded a total of 167 articles (Figure 2). Of these, 133 articles were found in PubMed, Hinari and Google Scholar and the remaining 34 were found from the desk review. After reviewing the titles and abstracts we excluded 143 articles due to irrelevance. The full text of the remaining 24 articles were downloaded and assessed for quality and relevance. An article from Sierra Leone was exclude because the outcome was not clearly stated (30); three articles from Zambia were excluded due to duplication (31), lack of clarity regarding outcome (32) or irrelevance (33); three articles from Nigeria (34), Togo (35) and Benin (35) were excluded due to low quality scores; and one article from Ethiopia was excluded because the full text was not available (36). The remaining 16 studies were included in the analysis (**Figure 2)**. The 16 articles reported on six cross-sectional studies from Ethiopia (14–17, 37, 38), one cross-sectional study from Kenya (13), five from Nigeria (39–43), two from Rwanda (44, 45), one article from Sudan (46) and one from Zambia (47).

**Figure 2.**
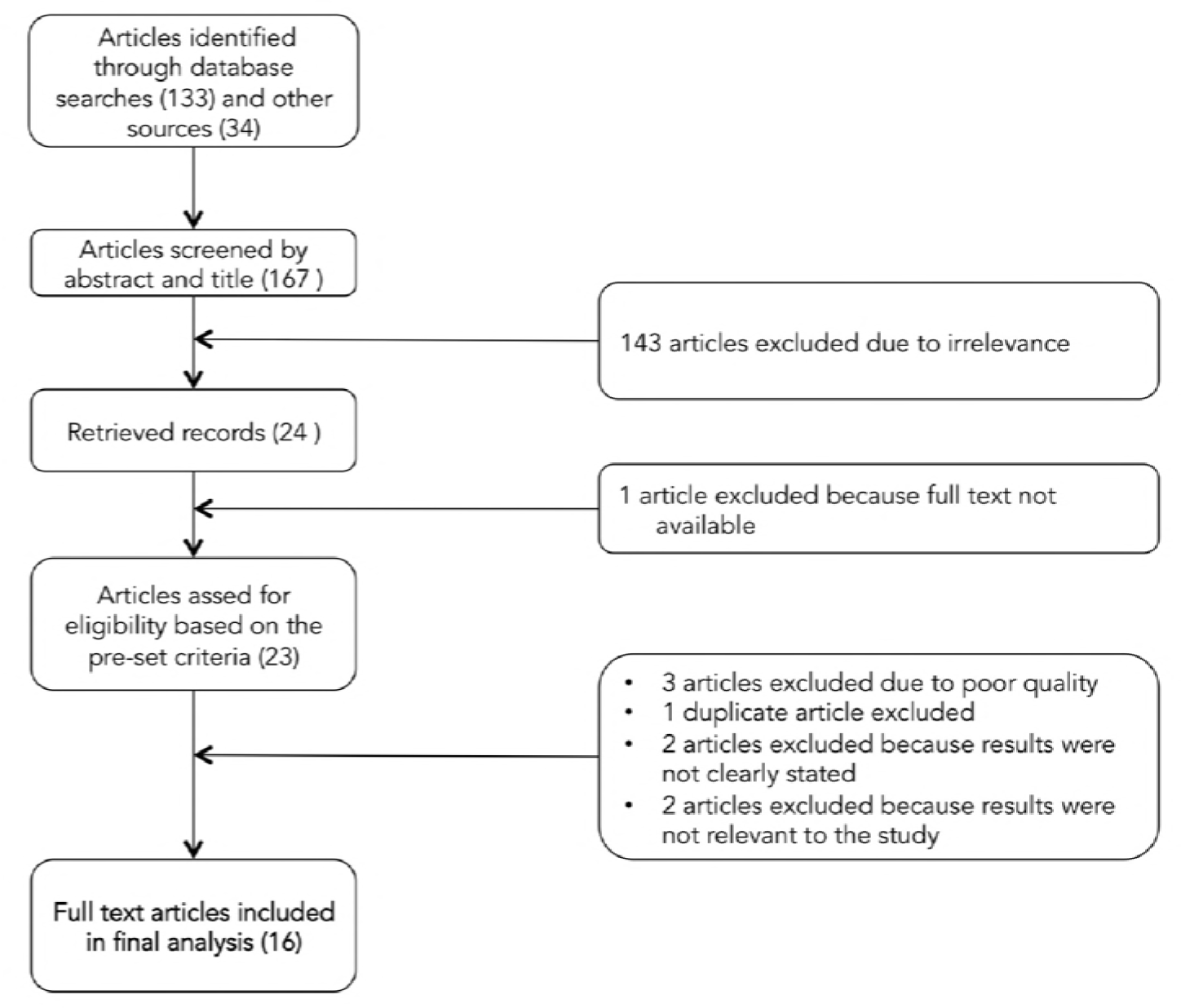
Flow diagram showing the procedure of selecting studies for meta-analysis, 2005–2017, sub-Saharan Africa.

### Characteristics of included studies

Sixteen studies with a total sample of 4,314 epileptic patients were assessed (Table 1). The majority of studies were conducted in Ethiopia (14–17, 37, 38), and Nigeria (39–43). The remaining were from Rwanda (44, 45), Sudan (46), Zambia (47) and Kenya (13). Studies were mostly reported in small, regional, peer-reviewed journals, with two reported in PhD dissertations.

**Table 1:** Characteristics of included studies for systematic review and meta-analysis, 2005–2017, sub-Saharan Africa.

| Study | Source Type | Study year | Country (Region) | Study design | Sample size | Response rate | Depression measure | Depression prevalence (%) | Quality Score | Factors positively associated with depression |
| --- | --- | --- | --- | --- | --- | --- | --- | --- | --- | --- |
| Tsegabrhan, et al (16) | Journal | 2014 | Ethiopia (East) | Cross-sectional | 300 | 100% | BDI | 49.3 | 7 | Lower educational status<br>Higher seizure frequency<br>Higher perceived stigma |
| Tegegne et al (15) | Journal | 2015 | Ethiopia (East) | Cross-sectional | 415 | 98% | HADS | 32.8 | 8 | Lower educational status<br>Polytherapy<br>Higher perceived stigma |
| Biftu et al (14) | Journal | 2015 | Ethiopia (East) | Cross-sectional | 405 | 96% | BDI | 45.2 | 7 | Lower educational status<br>Higher seizure frequency<br>Polytherapy<br>Higher perceived stigma<br>Early onset of seizures<br>Difficulty adhering to AEDs |
| Tegegne et al (38) | Journal | 2014 | Ethiopia (East) | Cross-sectional | 415 | 98% | HADS | 32.8 | 8 | (not assessed) |
| Biftu et al (37) | Journal | 2015 | Ethiopia (East) | Cross-sectional | 408 | 97% | BDI-II | 45.1 | 8 | (not assessed) |
| Tilahune et al (17) | Journal | 2016 | Ethiopia (East) | Cross-sectional | 326 | 100% | PHQ | 24.5 | 7 | Female gender<br>Greater age<br>Divorced<br>Lower income |
| Kiko (13) | Dissertation | 2013 | Kenya (East) | Cross-sectional | 327 | Not stated | BDI | 16.5 | 7 | Polytherapy<br>(associated with mild depression) |
| Adewuya et al (39) | Journal | 2005 | Nigeria (West) | Cross-sectional | 102 | 90% | DISC-IV | 28.4 | 8 | Polytherapy<br>Higher perceived stigma<br>Uncontrolled seizures |
| Ayanda et al (40) | Journal | 2016 | Nigeria (West) | Cross-sectional | 74 | Not stated | MINI | 21.6 | 8 | Not assessed |
| Owolabi et al (43) | Journal | 2016 | Nigeria (West) | Cross-sectional | 255 | Not stated | MINI | 20.4 | 8 | Higher seizure frequency<br>Early onset of seizures (duration of epilepsy)<br>Previous hospitalization for epilepsy |
| Mosaku et al (41) | Journal | 2006 | Nigeria (West) | Cross-sectional | 51 | Not stated | HADS | 27.5 | 7 | Not assessed |
| Ogunrin et al (42) | Journal | 2010 | Nigeria (West) | Case-control | 152 | Not stated | BDI, HRSD | 42 | 8 | Female gender<br>Uncontrolled seizures<br>Duration of epilepsy<br>Difficulty adhering to AEDs<br>(Associated with both BDI & HRSD) |
| Sezibera et al (45) | Journal | 2013 | Rwanda (East) | Cross-sectional | 105 | Not stated | HRSD | 48.6 | 8 | Female gender<br>Greater age<br>Lower educational status |
| Mutabazi (44) | Dissertation | 2014 | Rwanda (East) | Cross-sectional | 382 | Not stated | MINI | 6.5 | 8 | Not assessed |
| Saadalla et al (46) | Journal | 2016 | Sudan (East) | Case-control | 200 | Not stated | BDI | 45.5 | 7 | Not assessed |
| Veneviv et al (47) | Journal | 2016 | Zambia (South) | Cross-sectional | 397 | Not stated | BPRS | 39.4 | 7 | Not assessed |

Beck's Depression Inventory (BDI); Brief Psychiatric Rating Scale (BPRS); Diagnostic Interview Schedule for Children Version IV (DISC-IV); Hospital Anxiety and Depression Scale (HADS); Hamilton Rating Scale for Depression (HRSD); Mini International Neuropsychiatric Interview (MINI); Patient Health Questionnaire.

### Quality Assessment

Most studies were cross-sectional—two (42, 46) used a case-control design; and all recruited participants from in-patient or out-patient clinical settings.

Most studies had moderate sample sizes; only two had small samples of less than 100 participants (40, 41). Reported response rates were high (>90%) but more than half of the studies did not report a response rate and only one discussed the characteristics of non-responders. All studies used standardized methods for measuring depression with the Beck’s Depression Inventory (BDI) and the Hospital Anxiety and Depression Scale (HADS) being the tools most frequently used.

Half of the studies examined the factors associated with depression in PWE; three studies examined the factors related to PWE having any psychotic disorder or with them have a lower quality of life.

### Publication Bias

Both funnel plots of precision asymmetry and the Egger’s test of the intercept indicated the presence of publication bias in the studies. Visual examination of the funnel plot showed it to be asymmetric and Egger’s test of the intercept (B0) was 0.54 (95% CI: 0.2 - 0.87 p<0.05). To mitigate against publication bias we applied a trim and fill analysis in the random effects model; however, the prevalence estimates did not differ significantly between the initial model and the trim and fill model.

### Prevalence of depression in PWE

Depression prevalence ranged from 49.3% in an Ethiopian study (16) to 6.5%, reported in study conducted in Rwanda (44).

Because the I^2^ test for heterogeneity indicated significant difference between the studies (I^2^ =98 %, p<0.05) and because theoretically we expected that the study settings and socio-economic contexts might differ radically across these studies we fitted a DerSimonian and Laird random effect model to estimate the pooled prevalence of depression (48, 49). In the model each individual study is given a weight based its reported effect size and sample size (50). The studies with the largest weight were Michel (44), Kiko (13), and Tegegne et al (15) with respective weights of 6.5%, 6.43%, and 6.4%. Smaller weights were given for Mosaku, 5.65%, (41), Sezibera et al 5.97%, (45) and Ayanda et al 5.99% (40) (see Figure 3).

**Fig 3:**
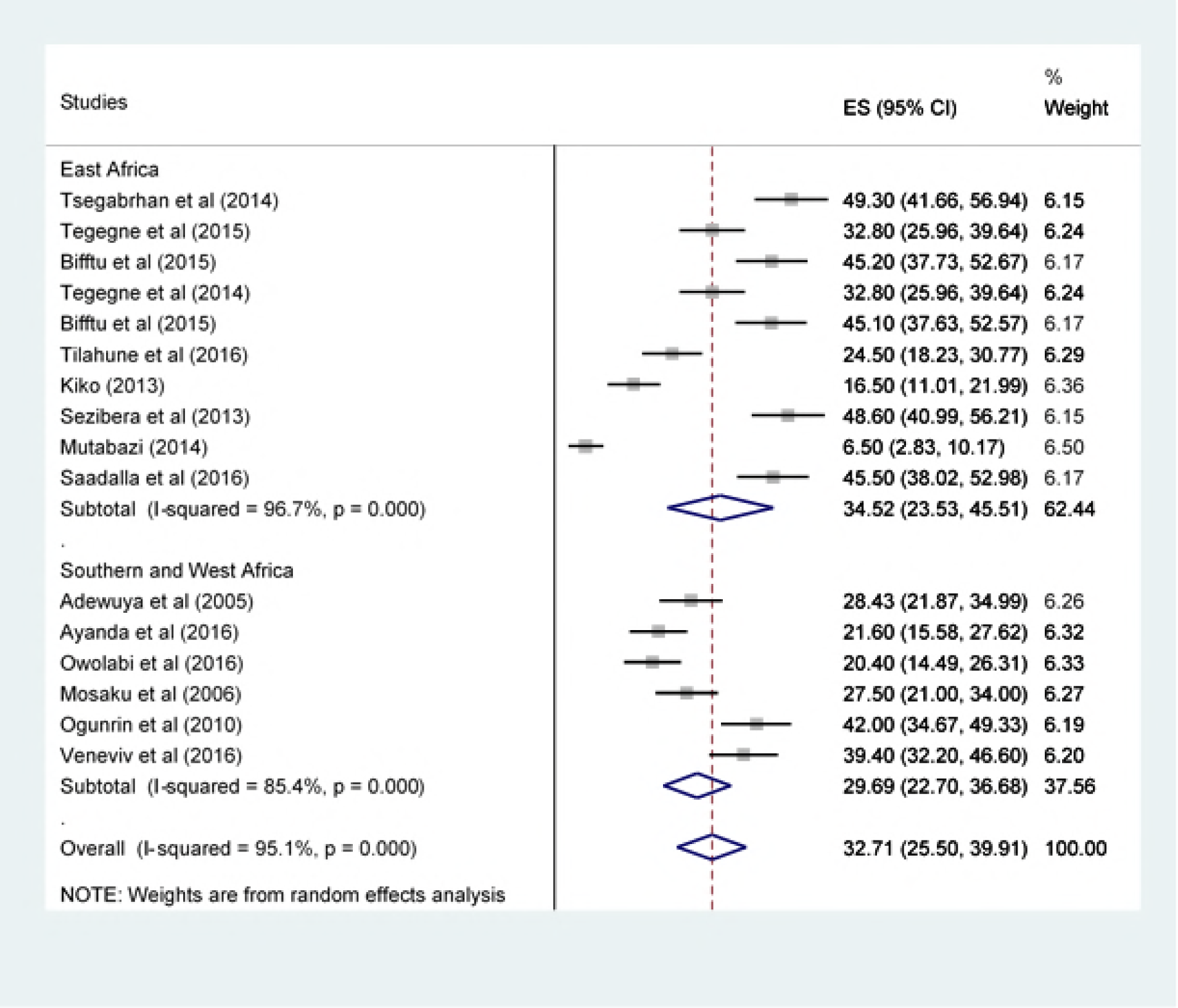
Forest plot of 16 studies assessing prevalence of depression among epilepsy patients, 2005–2017, sub-Saharan Africa.

The average pooled estimate of depression among epilepsy patients was 32.71 (95% CI: 25.50 - 39.91) (Figure 3). Sub-group analysis by geographic region found that the pooled prevalence of depression among epileptic patients in East Africa was 34.52 (95% CI: 23.53 - 45.51) and 29.69 (95% CI: 22.7 - 36.68) among patients in Southern and West Africa (Figure 3).

### Factors associated with depression among PWE

The factors most frequently associated with depression in PWE were, in order, lower educational status(14–16, 45), higher perceived stigma(14–16, 39), polytherapy(13, 15, 16, 39), female gender(17, 42, 45), the frequency of seizures(14, 16, 43) or having controlled seizures(39, 42), the duration of epilepsy(14, 42) and greater age(17, 45).

four of the studies reviewed explicitly intended to find factors associated with depression in epilepsy patients(14–16, 39). One of the factors most strongly associated with depression among PWE was and the amount of different medications patients received. The pooled odds of depression among epilepsy patients receiving polytherapy was 2.65 (95% CI: 1.49 - 4.71) compared with patients receiving monotherapy (Figure 4).

**Figure 4.**
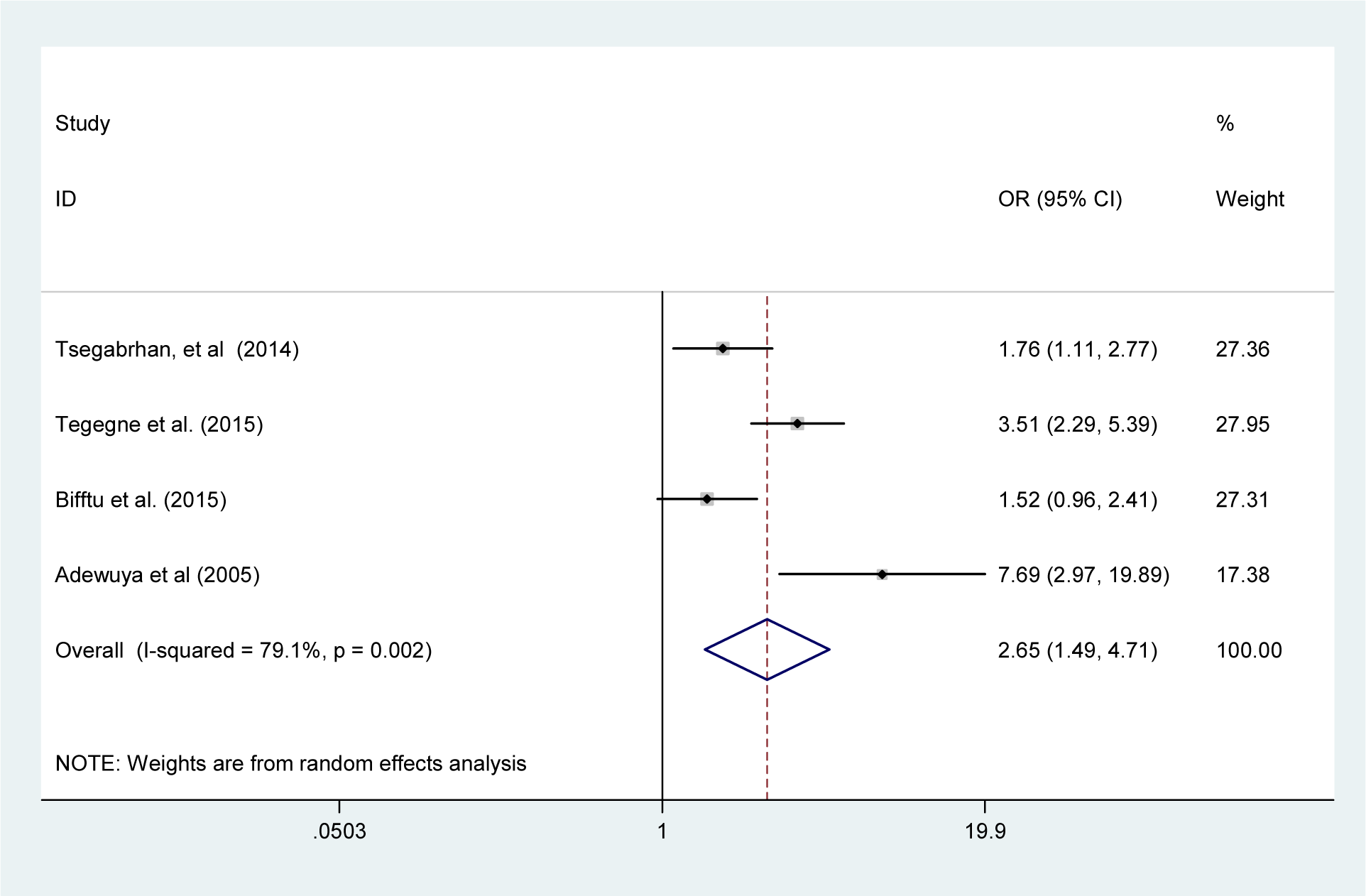
Forest plot of four studies examining the effect of polytherapy on depression among epilepsy patients 2005–2017, sub-Saharan Africa.

## Discussion

### Key Findings

This systemic review and meta-analysis attempted to estimate the pooled prevalence of depression and its association with polytherapy among epilepsy patients in sub-Saharan Africa. We found a very high prevalence of depression (32.71%) among PWE that was roughly 10 percentage points larger than that reported in recent meta-analyses; which reviewed studies that were almost exclusively conducted in Europe, the Americas, or Asia (21–23). In the one recent meta-analysis that included sub-Saharan Africa, the region was also found to have the highest regional prevalence of comorbidity although that pooled prevalence, 25.6% was lower than that found in our study (23).

While we expected to find significant comorbidity due to the well-established bi-directional relationship between the depression and epilepsy (2, 51), the high prevalence of depression reported in the studies under review suggest that the psychological toll of the epilepsy may be particularly severe in sub-Saharan African settings where the illness is more poorly managed, treatment options limited, epilepsy-related stigma high, and the social and economic costs of illness particularly acute. This is because the relationship between depression and epilepsy is not only neurological but also triggered and shaped by socioeconomic factors (52). In settings where epilepsy is thought to be a potentially contagious spiritual curse, PWE face profound social isolation and limited prospects for education, employment, or marriage (20) leaving them physically, emotionally, and economically vulnerable. In addition, endemic poverty in many communities means that PWE must manage the stress of affording treatment while maintaining their livelihoods in households that are often food insecure and financially precarious. The argument that high social stigma, poor treatment, and low socio-economic status in sub-Saharan Africa are important drivers of its higher rates of comorbidity is supported by the findings in the individual studies reviewed that these factors are significantly associated with depression among PWE.

A secondary objective of this study was to determine the effect of polytherapy on the prevalence of depression in PWE. We found that polytherapy doubled the odds of depression (OR= 2.65; 95% CI: 1.49 - 4.71) compared with monotherapy. Polytheraphy not only increases pill burden, making adherence, and therefore control of seizures, more difficult, but also increases the risk of adverse drug reactions and the severity of those reactions (53, 54). In addition patients taking multiple medications have higher risk of drug-to-drug interactions and may more prone to medication errors (53, 54). Our findings suggest that greater emphasis should be placed on assessing pill burden, tolerability, and potential drug interactions as well as on providing appropriate health education about drug regimens when designing epilepsy treatment guidelines and provider training curricula.

Our regional sub-group analysis found higher prevalence of depression in East Africa than in Southern and West Africa (34.52 % vs 29.69%). This difference may be due to the large number of East African studies that were conducted in Ethiopia, which has a particularly weak mental health care system compared to other regions. Poorly controlled epilepsy is a risk factor for depression and Ethiopia lacks the person power and infrastructure to manage epilepsy well for the majority of patients in need. For example, for a country of almost 100 million people, Ethiopia has only approximately 63 psychiatrists, 150 masters-level mental health professionals, and 200 bachelors-level psychiatry nurses (55) and most of these providers are concentrated in the capital city. Non-mental health professionals are often assigned to provide mental health services in Ethiopia, but studies indicate that they lack the knowledge required to deliver comprehensive mental health care (56). Ethiopia’s mental health system also faces many other challenges that complicate treatment of epilepsy such as delayed and inadequate supportive supervisions for trainees, lack of funding for supportive supervision and mentoring, and interrupted drug supplies (55). Ethiopia’s overall poverty compared to Zambia and Nigeria may also contribute to the difference in regional prevalence rates, since as discussed above, poor socio-economic status is associated with depression in PWE.

### Study Limitations

There are several limitations in this review. Firstly, we were only able to review English-language studies, which may have caused us to omit important studies from Francophone and Lusophone Africa. Secondly, as noted above, many of the studies reviewed were published in small regional journals or were PhD dissertations making it difficult to gauge the extent of peer review. Moreover, due to the absence of data, crude odds ratios were used to estimate factors related with comorbidity, which prevented us excluding confounding factors. The findings of this meta-analysis would be best interpreted keeping these analytical limitations and the limitations of the original studies in mind.

### Conclusion

This meta-analysis found that the prevalence of comorbid depression with epilepsy in sub-Saharan Africa was high, and of greater magnitude than that reported in other geographic regions. We also find that comorbidity is and significantly associated with polytherapy. Given the high overall prevalence of epilepsy in sub-Saharan Africa, more attention should be paid to increasing health education on epilepsy to reduce stigma; to incorporating depression screening and treatment into existing epilepsy programs; and to revising treatment guidelines on co-morbid depression to reduce polytherapy. Future research in sub-Saharan Africa should focus on identifying appropriate medication regimens for patients with comorbid depression.

## Abbreviations

AEDs: Anti-epileptic drugs
BDI: Beck’s Depression Inventory
BPRS: Brief Psychiatric Rating Scale
CI: Confidence interval
CT: Cheru Tesema
DISC-IV: Diagnostic Interview Schedule for Children Version IV
FW: Fasil Wagnew
GD: Getenet Dessie
HADs: Hospital Anxiety and Depression Scale
HRSD: Hamilton Rating Scale for Depression
HM: Henok Mulugeta
MINI: Mini International Neuropsychiatric Interview
NOS: Newcastle-Ottawa Scale
OR: Odds ratio
PHQ: Patient Health Questionnaire
PWE: People with epilepsy
SB: Sahai Burrowes
WHO: World Health Organization

## Declarations

### Ethics approval and consent to participate

Not applicable

### Consent for publication

Not applicable

### Availability of data and material

Data are available upon request to the corresponding author.

### Author contributions

FW conceived and designed the research protocol. GD and FW conducted the literature review, data extraction, and data analysis tasks, and led the interpretation and drafting of the manuscript. HM and CT assisted with quality control. SB assisted in data interpretation and manuscript preparation and language editing. All the authors (GD, FW, HM, CT, and SB) were involved in revising the manuscript and editing the manuscript.

## Acknowledgment

Not applicable

## Reference

1. World health organization. Epilepsy in the WHO African region. Bridging the Gap, Paswerk Bedrijven, Hoofdorp, Netherlands. 2004.

2. Kanner AM. Depression and epilepsy: a new perspective on two closely related disorders. Epilepsy currents. 2006;6(5):141–6.

3. Mula M, Schmitz B. Depression in epilepsy: mechanisms and therapeutic approach. Therapeutic advances in neurological disorders. 2009;2(5):337–44.

4. Jones JE, Hermann BP, Barry JJ, Gilliam F, Kanner AM, Meador KJ. Clinical assessment of Axis I psychiatric morbidity in chronic epilepsy: a multicenter investigation. The Journal of neuropsychiatry and clinical neurosciences. 2005;17(2):172–9.

5. Trivedi MH, Kurian BT. Managing depressive disorders in patients with epilepsy. Psychiatry (Edgmont). 2007;4(1):26.

6. Cramer JA, Blum D, Reed M, Fanning K. The influence of comorbid depression on seizure severity. Epilepsia. 2003;44(12):1578–84.

7. Cramer JA, Blum D, Reed M, Fanning K. The influence of comorbid depression on quality of life for people with epilepsy. Epilepsy & Behavior. 2003;4(5):515–21.

8. Yang Y, Gao X, Xu Y. The dilemma of treatments for epileptic patients with depression. International Journal of Neuroscience. 2015;125(8):566–77.

9. Noe KH, Locke DE, Sirven JI. Treatment of depression in patients with epilepsy. Current treatment options in neurology. 2011;13(4):371–9.

10. Cramer JA, Blum D, Fanning K, Reed M. The impact of comorbid depression on health resource utilization in a community sample of people with epilepsy. Epilepsy & Behavior. 2004;5(3):337–42.

11. Ettinger AB, Good MB, Manjunath R, Faught RE, Bancroft T. The relationship of depression to antiepileptic drug adherence and quality of life in epilepsy. Epilepsy & Behavior. 2014;36:138–43.

12. Hitiris N, Mohanraj R, Norrie J, Sills GJ, Brodie MJ. Predictors of pharmacoresistant epilepsy. Epilepsy research. 2007;75(2-3):192–6.

13. Kiko N. Prevalence and factors associated with depression among patients with epilepsy in a Kenyan tertiary care hospital 2013.

14. Bifftu BB, Dachew BA, Tiruneh BT, Tebeje NB. Depression among people with epilepsy in Northwest Ethiopia: a cross-sectional institution based study. BMC research notes. 2015;8(1):585.

15. Tegegne MT, Mossie TB, Awoke AA, Assaye AM, Gebrie BT, Eshetu DA. Depression and anxiety disorder among epileptic people at Amanuel Specialized Mental Hospital, Addis Ababa, Ethiopia. BMC psychiatry. 2015;15(1):210.

16. Tsegabrhan H, Negash A, Tesfay K, Abera M. Co-morbidity of depression and epilepsy in Jimma University specialized hospital, Southwest Ethiopia. Neurology India. 2014;62(6):649.

17. Tilahune AB, Bekele G, Mekonnen N, Tamiru E. Prevalence of unrecognized depression and associated factors among patients attending medical outpatient department in Adare Hospital, Hawassa, Ethiopia. Neuropsychiatric disease and treatment. 2016;12:2723.

18. Suljić E, Alajbegović A, Kucukalić A, Loncarević N. Comorbid depression in patients with epilepsy treated with single and multiple drug therapy. Medicinski arhiv. 2003;57(5-6 Suppl 1):45–6.

19. Ba-Diop A, Marin B, Druet-Cabanac M, Ngoungou EB, Newton CR, Preux P-M. Epidemiology, causes, and treatment of epilepsy in sub-Saharan Africa. The Lancet Neurology. 2014;13(10):1029–44.

20. Wilmshurst JM, Birbeck GL, Newton CR. Epilepsy is ubiquitous, but more devastating in the poorer regions of the world…or is it? Epilepsia. 2014;55(9):1322–5.

21. Scott AJ, Sharpe L, Hunt C, Gandy M. Anxiety and depressive disorders in people with epilepsy: A meta-analysis. Epilepsia. 2017;58(6):973–82.

22. Fiest KM, Dykeman J, Patten SB, Wiebe S, Kaplan GG, Maxwell CJ, et al. Depression in epilepsy a systematic review and meta-analysis. Neurology. 2013;80(6):590–9.

23. Kim M, Kim Y-S, Kim D-H, Yang T-W, Kwon O-Y. Major depressive disorder in epilepsy clinics: A meta-analysis. Epilepsy & Behavior. 2018;84:56–69.

24. Fdroemo H. National Mental Health Strategy 2012/13—2015/16. 2012.

25. Moher D, Liberati A, Tetzlaff J, Altman DG. Preferred reporting items for systematic reviews and meta-analyses: the PRISMA statement. Annals of internal medicine. 2009;151(4):264–9.

26. Stang A. Critical evaluation of the Newcastle-Ottawa scale for the assessment of the quality of nonrandomized studies in meta-analyses. European journal of epidemiology. 2010;25(9):603–5.

27. Huedo-Medina TB, Sánchez-Meca J, Marín-Martínez F, Botella J. Assessing heterogeneity in meta-analysis: Q statistic or I² index? Psychological methods. 2006;11(2):193.

28. Rendina-Gobioff G. Detecting publication bias in random effects meta-analysis: An empirical comparison of statistical methods. 2006.

29. Talebi M. Study of publication bias in meta-analysis using trim and fill method. International Research Journal of Applied and Basic Sciences. 2013;4(1):31–6.

30. M’bayo T, Tomek M, Kamara C, Lisk DR. Psychiatric comorbidity in African patients with epilepsy–Experience from Sierra Leone. International Journal of Epilepsy. 2017;4(1):26–30.

31. Mbewe EK, Uys LR, Birbeck GL. Detection and management of depression and/or anxiety for people with epilepsy in primary health care settings in Zambia. Seizure. 2013;22(5):401–2.

32. Mbewe EK. Improving Detection of Depression And/or Anxiety as Comorbidities of Epilepsy in Primary Health Care Settings in Zambia: University of KwaZulu-Natal, Durban; 2013.

33. Mbewe EK, Uys LR, Nkwanyana NM, Birbeck GL. A primary healthcare screening tool to identify depression and anxiety disorders among people with epilepsy in Zambia. Epilepsy & Behavior. 2013;27(2):296–300.

34. Onwuekwe I, Ekenze O, Bzeala-Adikaibe O, Ejekwu J. Depression in patients with epilepsy: a study from Enugu, South East Nigeria. Annals of medical and health sciences research. 2012;2(1):10–3.

35. Nubukpo P, Preux P, Houinato D, Radji A, Grunitzky E, Avode G, et al. Psychosocial issues in people with epilepsy in Togo and Benin (West Africa) I. Anxiety and depression measured using Goldberg’s scale. Epilepsy & Behavior. 2004;5(5):722–7.

36. Prevalence and associated factors of depression among epileptic patients attending outpatient department at hawassa university referral comprehensive hospital,Hawassa,SNNPR,Ethiopia 2017 G.C.

37. Bifftu BB, Dachew BA, Tiruneh BT. Perceived stigma and associated factors among people with epilepsy at Gondar University Hospital, Northwest Ethiopia: a cross-sectional institution based study. African health sciences. 2015;15(4):1211–9.

38. Tegegne MT, Muluneh NY, Wochamo TT, Awoke AA, Mossie TB, Yesigat MA. Assessment of quality of life and associated factors among people with epilepsy attending at Amanuel Mental Specialized Hospital, Addis Ababa, Ethiopia. Science Journal of Public Health. 2014;2(5):378–83.

39. Adewuya AO, Ola BA. Prevalence of and risk factors for anxiety and depressive disorders in Nigerian adolescents with epilepsy. Epilepsy & behavior. 2005;6(3):342–7.

40. Ayanda KA, Sulyman D. The predictors of psychiatric disorders among people living with epilepsy as seen in a Nigerian Tertiary Health Institution. Nigerian medical journal: journal of the Nigeria Medical Association. 2016;57(1):24.

41. Mosaku KS, Fatoye FO, Komolafe M, Lawal M, Ola BA. Quality of life and associated factors among adults with epilepsy in Nigeria. The International Journal of Psychiatry in Medicine. 2006;36(4):469–81.

42. Ogunrin OA, Obiabo YO. Depressive symptoms in patients with epilepsy: Analysis of self-rating and physician’s assessment. Neurology India. 2010;58(4):565.

43. Owolabi SD, Owolabi LF, Udofia O, Sale S. Depression in patients with epilepsy in Northwestern Nigeria: Prevalence and clinical correlates. Annals of African medicine. 2016;15(4):179.

44. Michel M. Adherence and Treatment Outcomes among Patients with Comorbidity of Depression and Other Mental Disorders attending Psychiatric Hospitals in Rwanda: Kenyatta University; 2014.

45. Sezibera V, Nyirasafari D. Incidence of depression in Epilepsy patients. Rwanda Journal. 2013;1(1):67–77.

46. Saadalla A, Elbadwi A. Depression among Sudanese epileptic patients. Age.5(5):18–25.

47. Venevivi L, Mbewe E, Paul R. Determining treatment levels of comorbid psychiatric conditions in people with epilepsy attending selected local clinics in Lusaka, Zambia. Medical Journal of Zambia. 2016;43(4):184–90.

48. Kelley GA, Kelley KS. Statistical models for meta-analysis: A brief tutorial. World journal of methodology. 2012;2(4):27.

49. Jackson D, Bowden J, Baker R. How does the DerSimonian and Laird procedure for random effects meta-analysis compare with its more efficient but harder to compute counterparts? Journal of Statistical Planning and Inference. 2010;140(4):961–70.

50. Higgins JP, Thompson SG, Deeks JJ, Altman DG. Measuring inconsistency in meta-analyses. BMJ: British Medical Journal. 2003;327(7414):557.

51. Hesdorffer DC, Hauser WA, Olafsson E, Ludvigsson P, Kjartansson O. Depression and suicide attempt as risk factors for incident unprovoked seizures. Annals of neurology. 2006;59(1):35–41.

52. Sankar R, Mazarati A. Neurobiology of depression as a comorbidity of epilepsy. Epilepsia. 2010;51(s5):81-.

53. Duryea PB, Moore C, Nathanson-Shinn A, Hall SE. Psychiatric Polypharmacy: A Word of Caution.

54. Kukreja S, Kalra G, Shah N, Shrivastava A. Polypharmacy in psychiatry: a review. Mens sana monographs. 2013;11(1):82.

55. Ayano G, Assefa D, Haile K, Bekana L. Experiences, Strengths and Challenges of Integration of Mental Health into Primary Care in Ethiopia. Experiences of East African Country. Fam Med Med Sci Res. 2016;5(204):2.

56. Abera M, Tesfaye M, Belachew T, Hanlon C. Perceived challenges and opportunities arising from integration of mental health into primary care: a cross-sectional survey of primary health care workers in south-west Ethiopia. BMC health services research. 2014;14(1):113.

